## Supplementary Information for "Bayesian-Maximum-Entropy reweighting of IDP ensembles based on NMR chemical shifts"

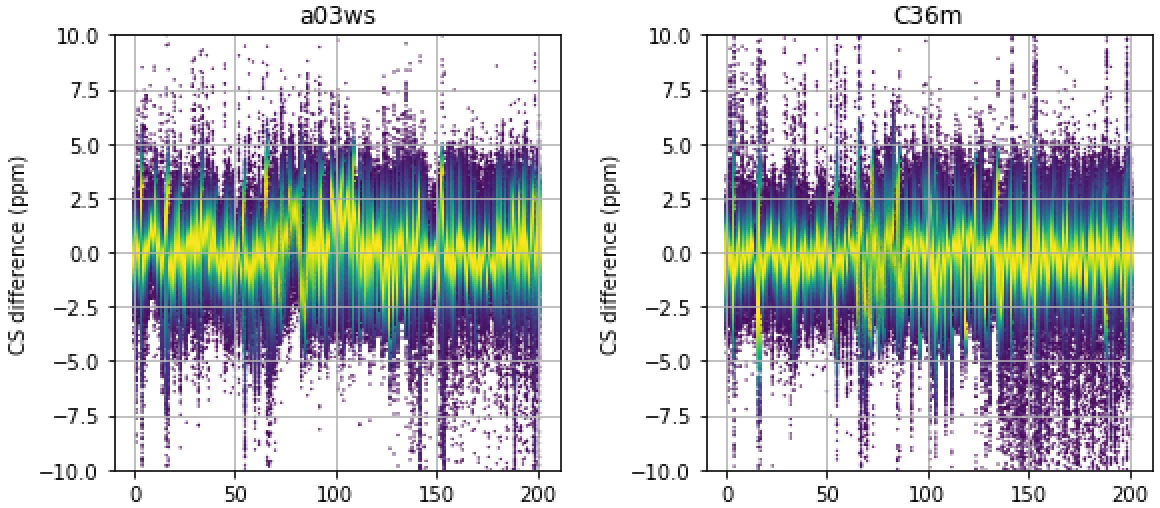


Figure S1. Distribution of the difference between the ensemble CS and the average target CS for the 2 ensembles discussed in this work. The x-values correspond to a concatenation of C, CA and CB CS. To be able to reweight the values should be distributed at both sides of y=0, as is the case.

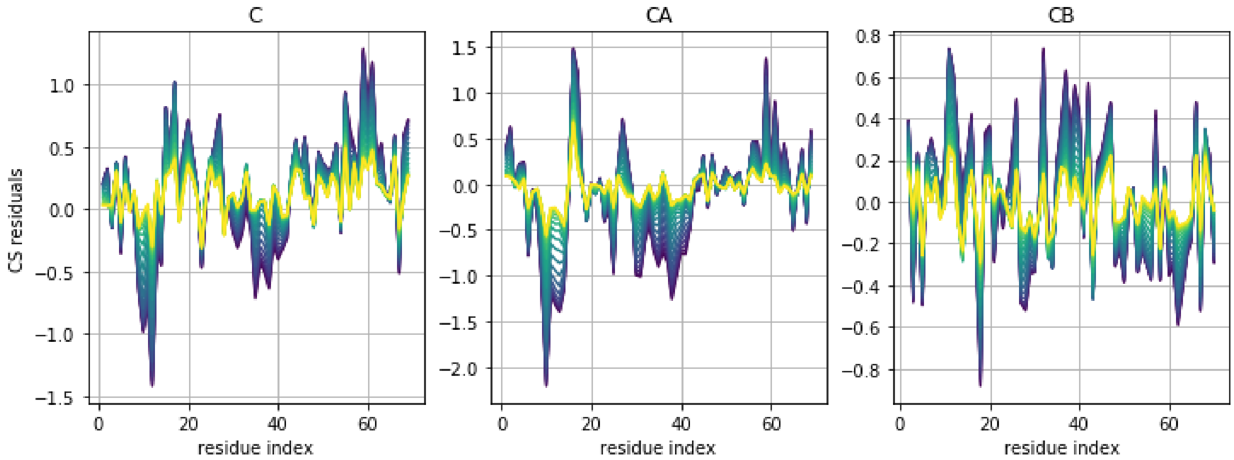


Figure S2. Evolution of the CS difference (residuals) for the reweighting procedure for a range of θ values from 100 (purple values) to 0.1 (yellow values). As the residuals have the same sign for most of the reweighting procedure, one cannot use the Wald-Wolfowitz run test to define the amount of reweighting.

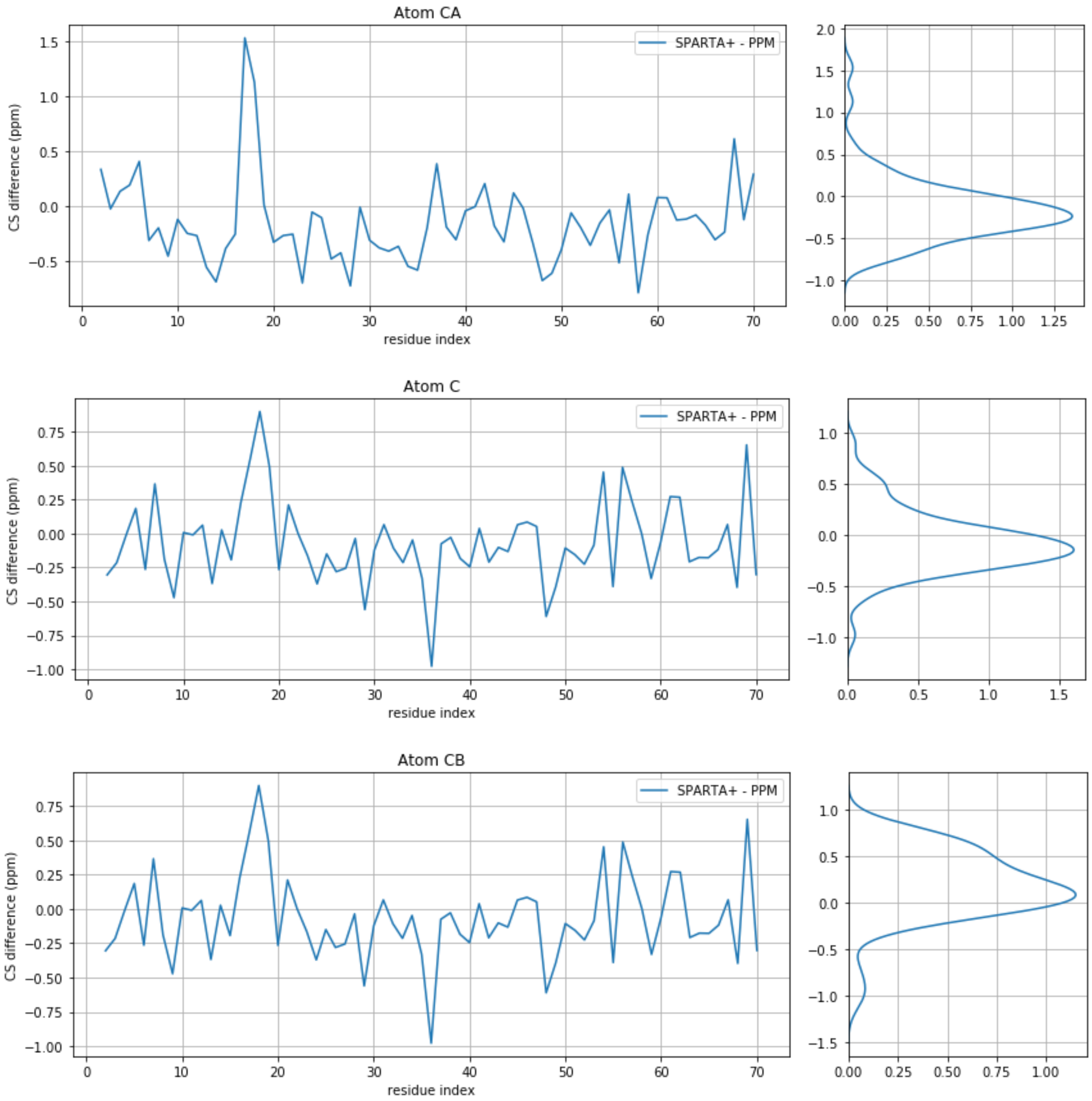


Figure S3. Difference in the chemical shifts predicted with Sparta+ and PPM for the a99SBdisp ensemble. The left plots show the value for each residue and the right plots show their distribution for a better visualization of their spread and shift. Atom types are CA (top), C (middle) and CB) bottom.

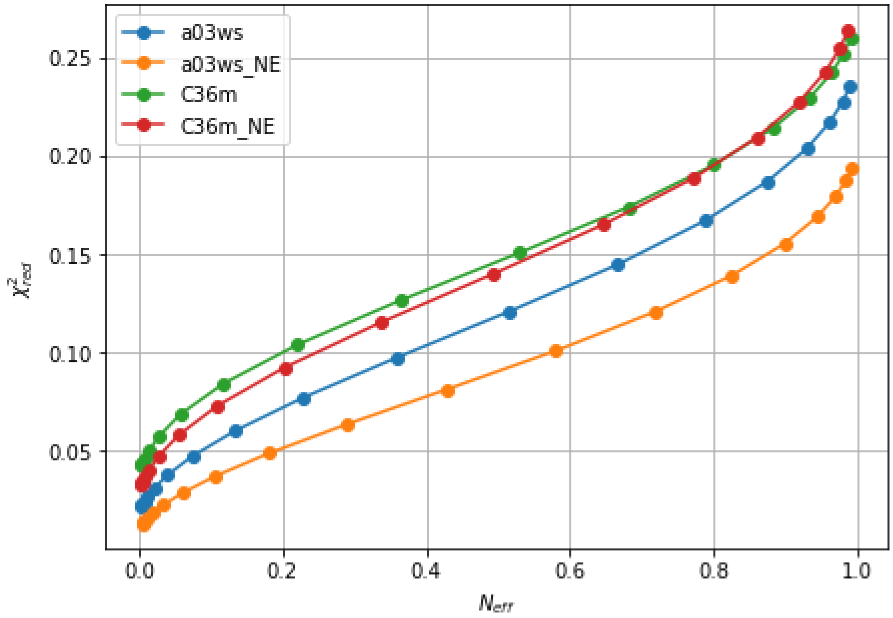


Figure S4. Evolution of $\chi^2_{red}$ with the effective sample size ($N_{eff}$), showing the lack of an L-shaped curve. The blue and orange lines are the same as in Figure 1. The NE suffix stands for “no-error”. It corresponds to the values where the target CS are calculated with PPM. For the sake of clarity $\chi^2_{red}$ uses the same error as in the case with errors, even though $\chi^2_{red}$ is ill-defined in this case and should be regarded as a scaled root-mean-square error.

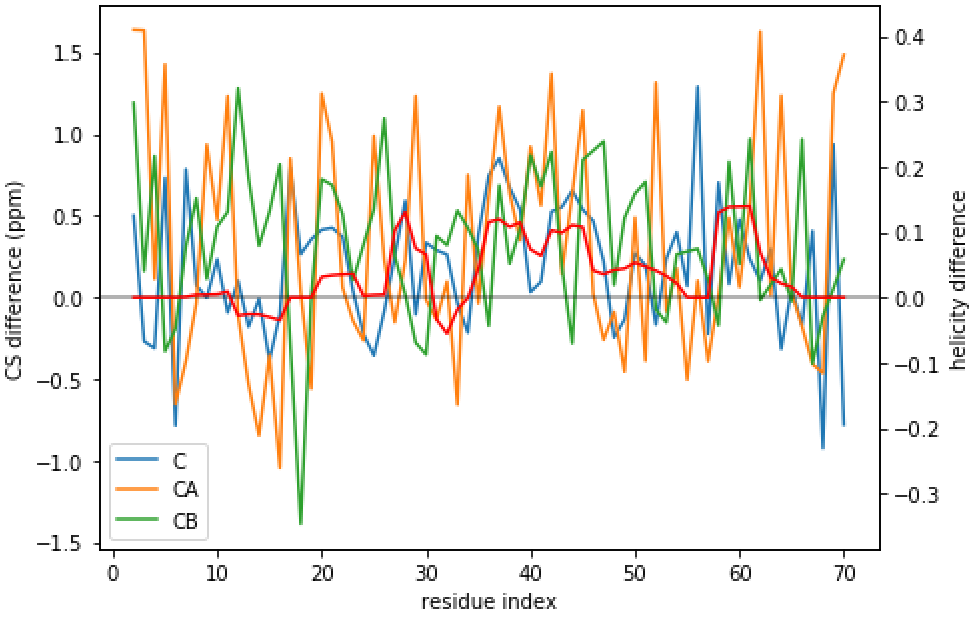


Figure S5. The CS difference between the target CS and the computed CS for the C36m ensemble compared to the difference in helicity for these two ensembles.

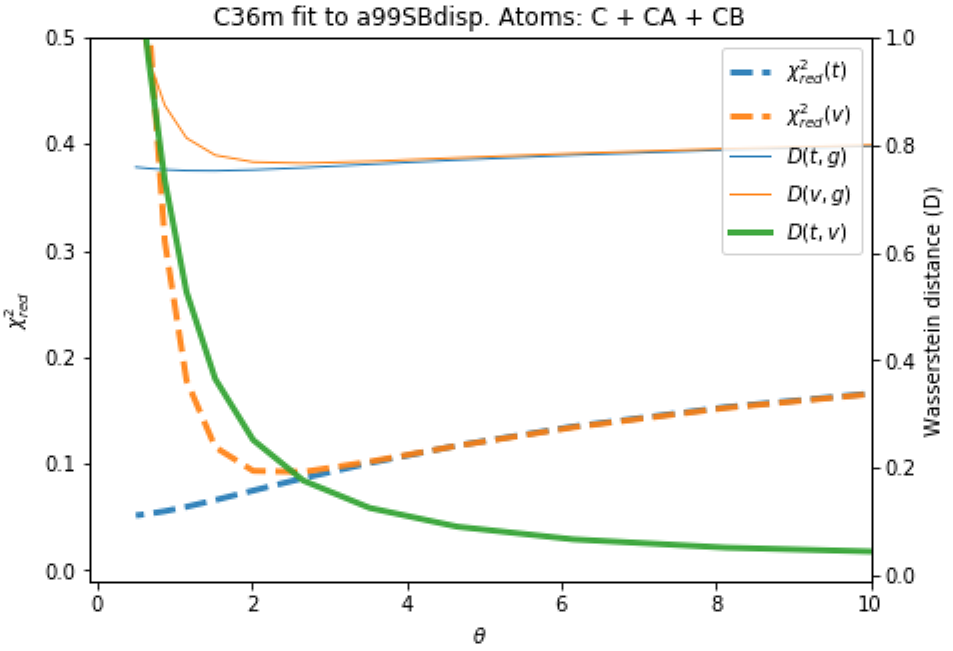


Figure S6. Behaviour of different quantities during the reweighting procedure of the C36m ensemble.

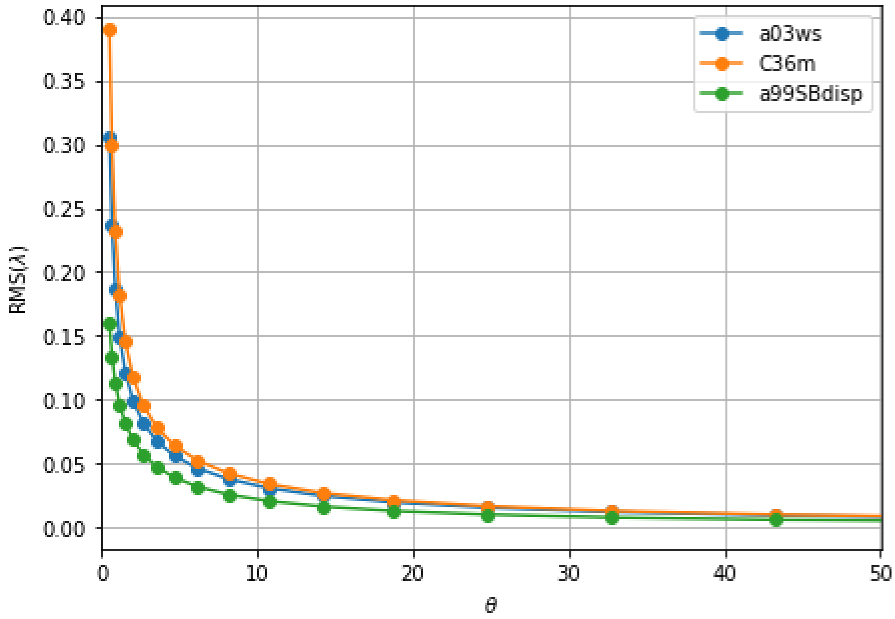


Figure S7. Root mean square of the Lagrange parameters λ during the reweighing procedure for all three ensembles. See Eq. 4.

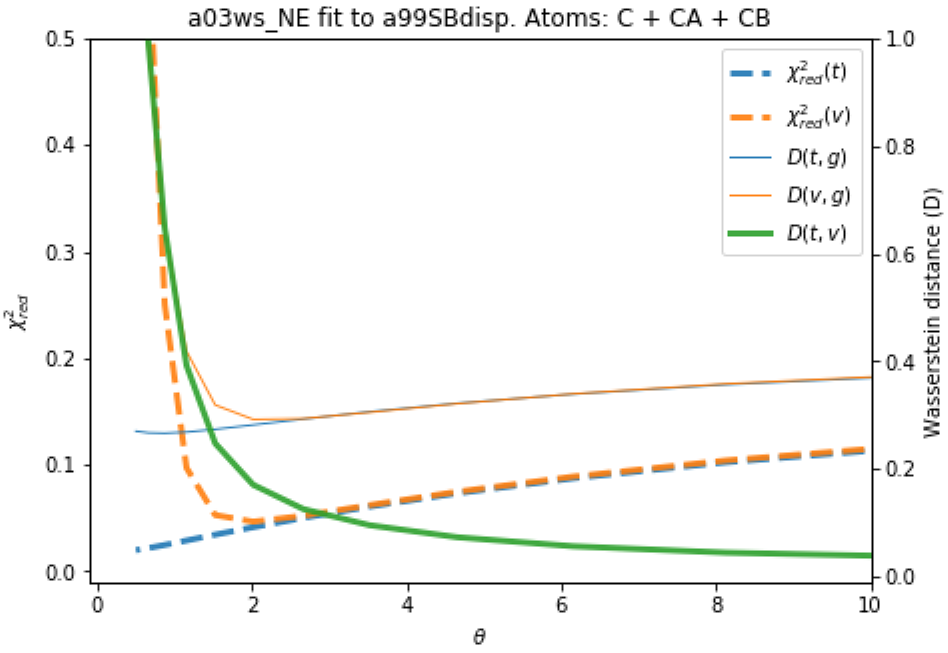


Figure S8. Behaviour of different quantities during the reweighting procedure of the a03ws ensemble with no predictive errors (NE suffix). As was done in figure S3, $\chi^2_{red}$ uses the same error as in the case with errors, even though $\chi^2_{red}$ is ill-defined in this case and should be regarded as a scaled root-mean-square error.

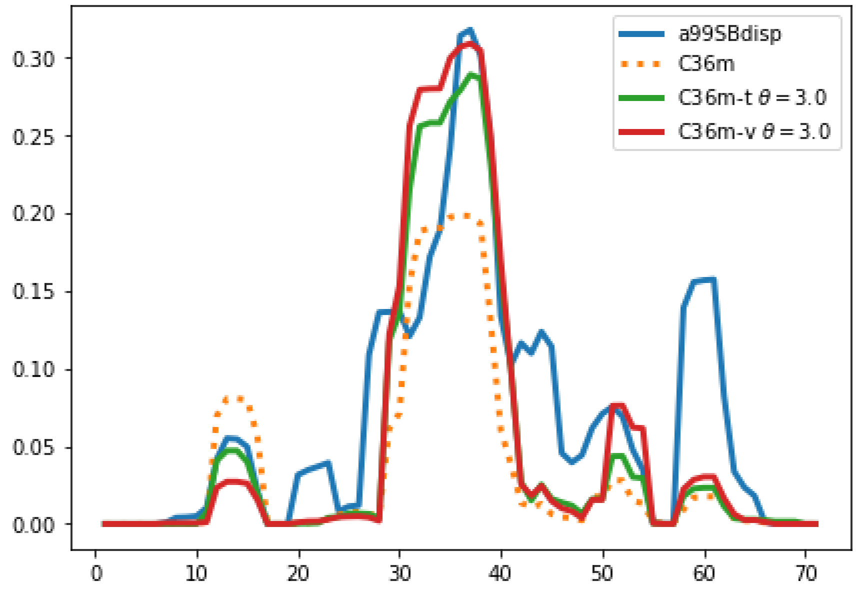


Figure S9. Alpha-helical content for the reweighted and target ensembles for the C36m force field. For the reweighted ensembles the train (C36m-t) and validation (C36m-v) sets are shown. The original ensembles before the reweighting is also shown.

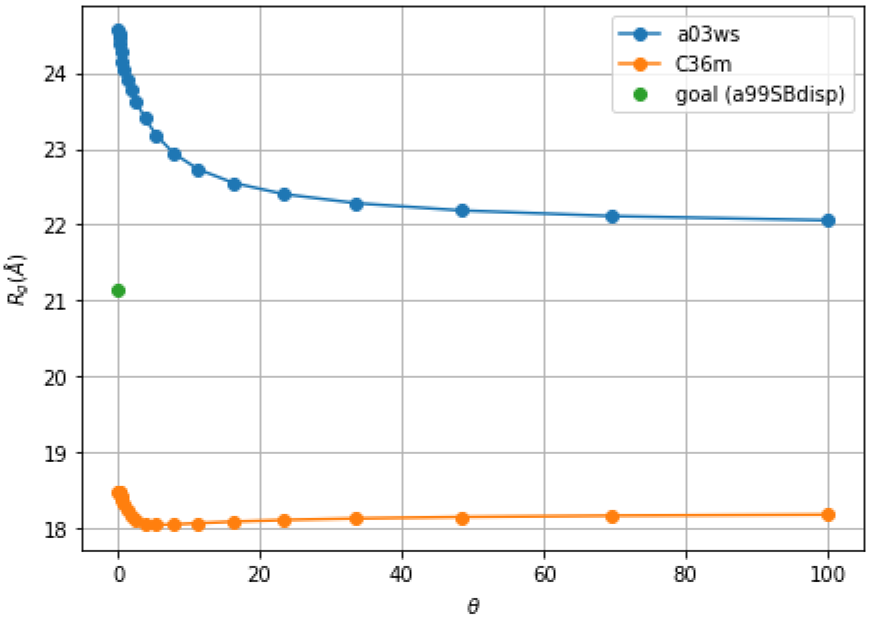


Figure S10. Evolution of the radius of gyration for different reweightings.

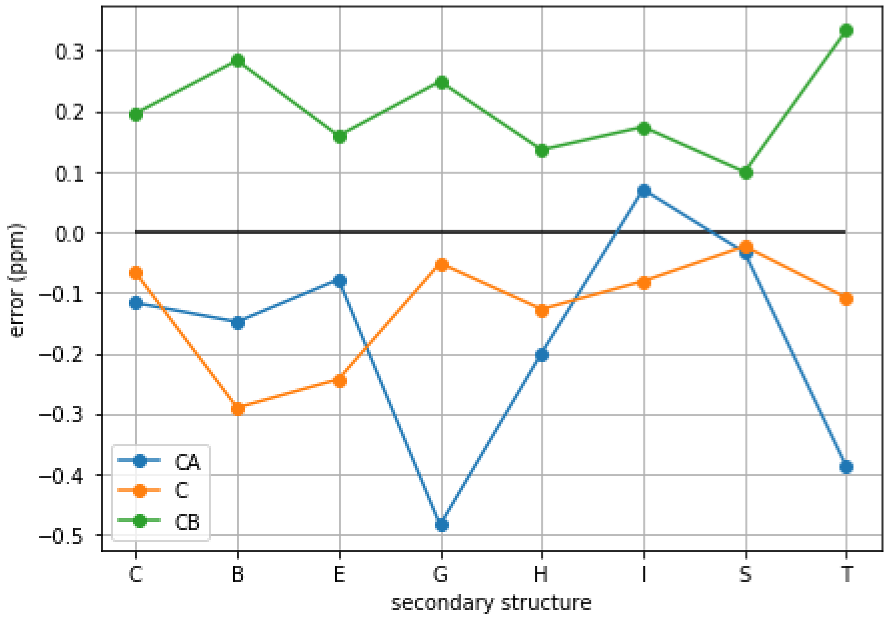


Figure S11. Error in the predictor (PPM) with respect to the target chemical shift (Sparta+) for different secondary structure elements of the a99SBdisp ensemble. For atoms CA and C there is a systematic underestimation of the chemical shift, whereas for CB, there is a systematic overestimation. The errors also depend on the type of secondary structure.  The codes are the following: ‘C‘ : Loops and irregular elements, ‘B’ : Residue in isolated beta-bridge, ‘E’ : Extended strand, participates in beta ladder, ‘G’ : 3-helix (3/10 helix), ‘H’ : Alpha helix, ‘I’ : 5 helix (pi helix), ‘S’ : bend, and ‘T’ : hydrogen bonded turn, as determined by the DSSP algorithm implemented in MDtraj.

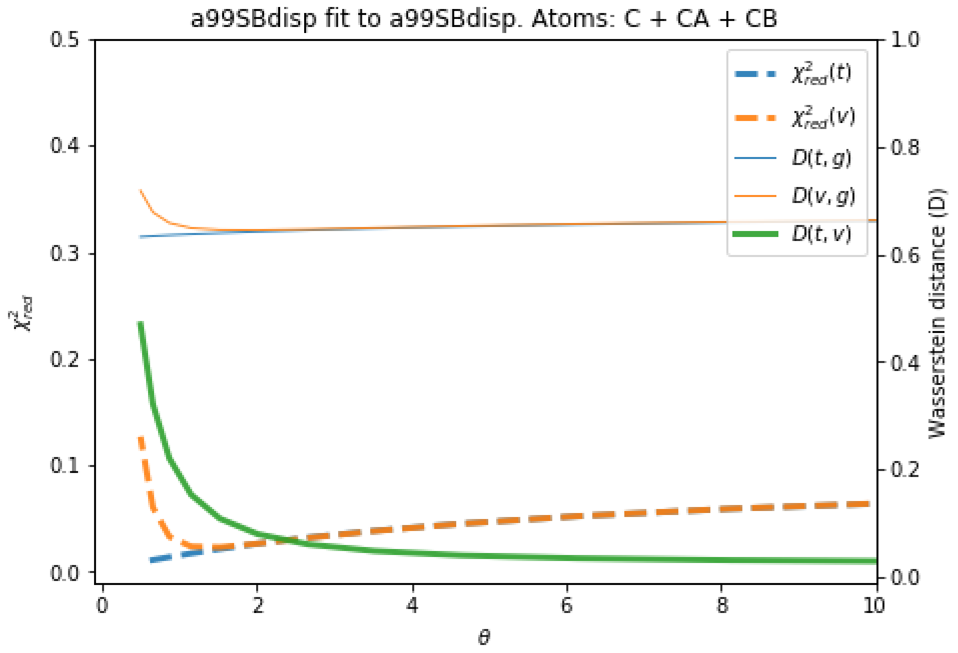


Figure S12. Behaviour of different quantities during the reweighting procedure of the a99SBdisp ensemble. The quantities defined as thin lines would not be measurable in a real case scenario, but their behaviour can be inferred from the quantities in thick lines.

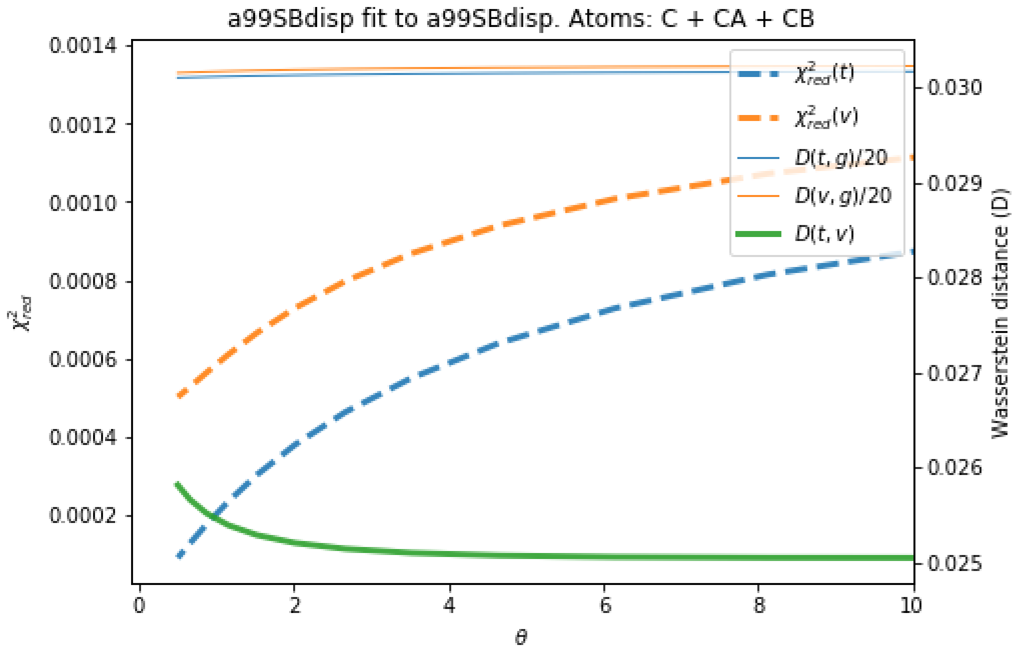


Figure S13. Behaviour of different quantities during the reweighting procedure of the a99SBdisp ensemble using secondary chemical shifts. The quantities defined as thin lines would not be measurable in a real case scenario, but their behaviour can be inferred from the quantities in thick lines. Remark that the $\chi^2_{red}$ values are very small for all θ values. *D(t,g)* and *D(t,v)* have been scaled by 1/20 so that the shape of the curves could be seen.

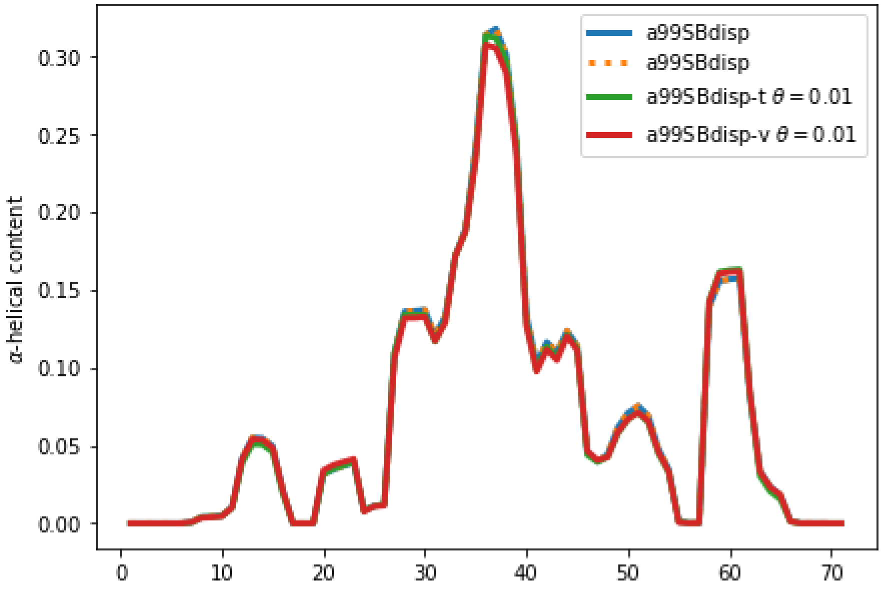


Figure S14. Alpha-helical content for the reweighted and target ensembles using secondary chemical shifts. For the reweighted ensembles the train (a99SBdisp-t) and validation (a99SBdisp-v) sets are shown. The original ensembles before the reweighting is also shown and, as expected, it corresponds exactly to the target ensemble as they are the same.

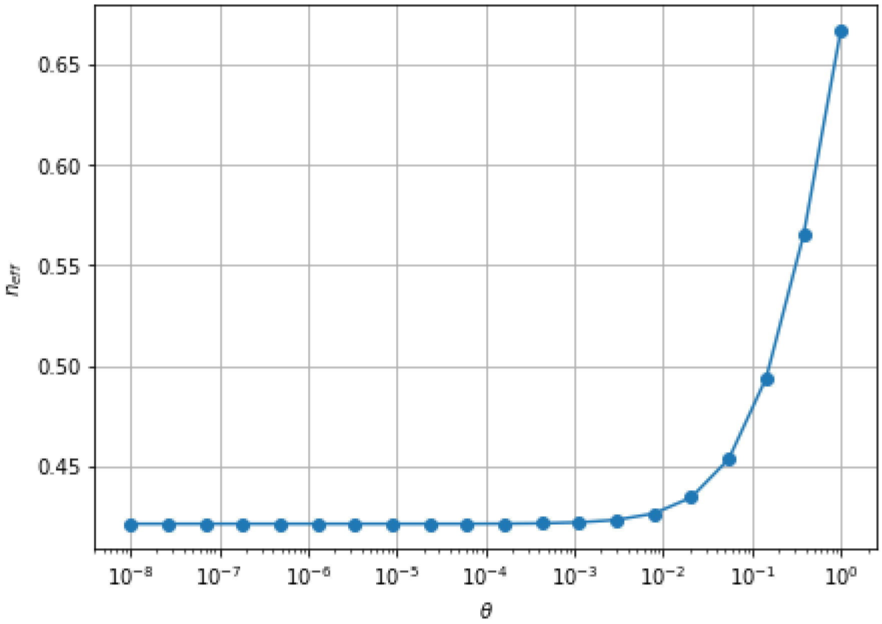


Figure S15. Evolution of the effective sample size $(N_{eff})$ with θ for the secondary chemical shift fitting of a99SBdisp ensemble to the a99SBdisp target ensemble.
